# Diagnostic algorithms to study post-concussion syndrome using electronic health records: validating a method to capture an important patient population

**DOI:** 10.1101/336560

**Authors:** Jessica Dennis, Aaron M. Yengo-Kahn, Paul Kirby, Gary S. Solomon, Nancy J. Cox, Scott L. Zuckerman

**Affiliations:** Division of Genetic Medicine, Department of Medicine, Vanderbilt University Medical Center Nashville, TN; Vanderbilt Genetics Institute, Vanderbilt University Medical Center, Nashville, TN; Vanderbilt Sports Concussion Center, Vanderbilt University School of Medicine, Nashville, TN; Department of Neurological Surgery, Vanderbilt University School of Medicine, Nashville, TN

**Keywords:** Post-concussion syndrome, electronic health records, diagnostic algorithm, genetics

## Abstract

**Introduction:** Post-concussion syndrome (PCS) is characterized by persistent cognitive, somatic, and emotional symptoms after a mild traumatic brain injury (mTBI). Genetic and other biological variables may contribute to PCS etiology, and the emergence of biobanks linked to electronic health records (EHR) offers new opportunities for research on PCS. We sought to validate the use of EHR data of PCS patients by comparing two diagnostic algorithms.

**Methods:** Vanderbilt University Medical Center curates a de-identified database of 2.8 million patient EHR. We developed two EHR-based algorithmic approaches that identified individuals with PCS by: (i) natural language processing (NLP) of narrative text in the EHR combined with structured demographic, diagnostic, and encounter data; or (ii) coded billing and procedure data. The predictive value of each algorithm was assessed, and cases and controls identified by each approach were compared on demographic and medical characteristics.

**Results:** First, the NLP algorithm identified 507 cases and 10,857 controls. The positive predictive value (PPV) in the cases was 82% and the negative predictive value in the controls was 78%. Second, the coded algorithm identified 1,142 patients with two or more PCS billing codes and had a PPV of 76%. Comparisons of PCS controls to both case groups recovered known epidemiology of PCS: cases were more likely than controls to be female and to have pre-morbid diagnoses of anxiety, migraine, and PTSD. In contrast, controls and cases were equally likely to have ADHD and learning disabilities, in accordance with the findings of recent systematic reviews of PCS risk factors.

**Conclusions:** EHR are a valuable research tool for PCS. Ascertainment based on coded data alone had a predictive value comparable to an NLP algorithm, recovered known PCS risk factors, and maximized the number of included patients.

---

More than 1.7 million Americans visit an emergency department or are hospitalized for a traumatic brain injury (TBI) each year, a number that underestimates the true burden due to those who do not seek medical care.^1^ TBI disproportionately affects the young and the old, and while motor vehicle collisions are a leading cause of TBI,^1, 2^ TBIs sustained in sport and military combat are increasingly recognized.^3, 4^

Up to 90% of TBIs are classified as mild (mTBI), defined as positive brain imaging with Glasgow Coma Scale (GCS) scores of 13-15, or concussion, defined as negative brain imaging with a similar GCS score range.^2, 5^ Symptoms of an mTBI may include temporary confusion, loss of consciousness, amnesia, or other neurological abnormality such as seizure,^6, 7^ and these symptoms typically resolve spontaneously or within seven to 10 days.^5, 8-10^ Up to 30% of patients, however, do not recover in this time and are unable to return to work, school, or other pre-injury activities for up to several months.^8, 11, 12^ These patients complain of persistent somatic, cognitive, and emotional symptoms, known clinically as post-concussion syndrome (PCS).

The determinants of PCS are poorly understood. Females and those with low educational attainment endure a longer recovery, as do patients with a personal and/or family history of headaches, migraines, and mood disorders.^11-16^ The literature is mixed, however, with regards to the effects of age, attention deficit hyperactive disorder (ADHD), and learning disability, though these factors may increase the risk of injury.^17, 18^ Initial injury severity and symptom burden also contribute to PCS risk, fueling the view that PCS is the result of both trauma to the brain and underlying psychological factors.^17, 19^

Despite this knowledge, PCS prediction tools explain at most 30% of the variability in mTBI outcome,^11-16^ spurring research into the role of other biological factors in recovery, including genetics.^20^ Preliminary studies in small selected samples have found associations between candidate genes and short- and long-term TBI outcomes, but aside from the *APOE e4* allele, which, in a meta-analysis of 14 studies^21^ increased the risk of poor 6 month outcomes by 36%, results have yet to be replicated in larger, representative populations. Moreover, there has been no agnostic search for genetic determinants of TBI recovery, as in a genome-wide association study (GWAS). Such hypothesis-free approaches have revealed novel biology in many rare and common diseases,^22^ but require very large numbers of cases and controls to be adequately powered, and replication of significant findings in an independent dataset to avoid false positives.^23^

Electronic health records (EHRs) linked to biorepositories such as DNA databanks offer new opportunities to study the biology of TBI recovery and PCS. EHRs are a rich data source, including information on pre-injury medical conditions, time-of-injury procedures, diagnoses, and post-injury follow-up care. The sample sizes afforded by EHRs are unprecedented, and since these data are collected in the course of routine clinical care, EHR studies are less costly and time-consuming than clinically recruited samples or prospective trials. EHR-linked biobanks are also growing in number and scale worldwide. Within the United States, 10 institutions have joined the National Human Genome Research Institute-funded Electronic Medical Records and Genomics (eMERGE) network (https://emerge.mc.vanderbilt.edu) since 2007, with at least 126,000 samples genotyped to date. More ambitiously, the National Institutes of Health Precision Medicine Initiative *All of US* program (https://allofus.nih.gov) aims to recruit and follow over one million Americans, linking their EHR to physical assessments, participant surveys, mobile health technologies, and biospecimens. Similar data are already available from the first 500,000 participants enrolled in the UK Biobank (http://www.ukbiobank.ac.uk) as well as the first 500,000 enrolled in the US Department of Veterans Affairs’ Million Veteran Program (http://www.research.va.gov/mvp), and studies using these data across a range of disorders and research questions are proving the value of EHR-recruited samples.^24-26^

Defining valid study populations is a necessary first step to using EHR data. Manual review of EHRs to identify a gold standard case-control set is time consuming, labor intensive, and impractical for large datasets. Algorithms are more efficient, and usually take one of two forms.^24^ Code-based algorithms apply filters to structured, coded data such as demographics, diagnostic and procedure codes, medications, and laboratory results. Algorithms based on natural language processing (NLP), on the other hand, use keyword searches and text mining software to extract data from unstructured text in the EHR. Both approaches have demonstrated merit,^24^ and NLP-based algorithms are typically used in conjunction with filters on coded data. An algorithm’s performance is often judged by its positive predictive value (PPV), the probability that a case identified by the algorithm is a true case, and its negative predictive value (NPV), the probability that a control identified by the algorithm is free of the disease under study.

Given the far-reaching public health implications of mTBI and the need for research, we sought to study mTBI recovery using large patient datasets. The objective of our study was to evaluate the validity of two algorithms for identifying PCS cases and controls from EHR data: (i) an NLP algorithm with additional filters applied to coded data; and (ii) a coded algorithm, based on only a limited set of filters applied to coded data. Our evaluation proceeded in two stages. First, we manually reviewed a subset of records identified by the NLP and coded algorithms and calculated positive and negative predictive values. Second, we compared all identified cases and controls on known PCS risk factors to determine the epidemiology of PCS in an EHR-recruited sample. We conclude with recommendations for designing future EHR algorithms to diagnose TBI and PCS.

## Methods

### Data Source

Vanderbilt University Medical Center (VUMC) is a tertiary care hospital in Nashville, Tennessee, with affiliated clinics throughout the region. EHRs have been used at VUMC since the early 1990s, and data are compiled in a de-identified IBM Netezza database known as the Synthetic Derivative (SD)^27^ The SD contains more than 2.8 million patient EHRs, and these are linked to DNA biosamples in >275,000 patients, who could eventually be used in genetic studies of PCS. Available data in the SD include basic demographics, billing codes, procedure codes, clinical documents, medications, laboratory values, and dates of inpatient stays and outpatient visits. The majority of billing codes are International Classification of Diseases, Ninth Revision (ICD-9), with VUMC transitioning to Tenth Revision (ICD-10) in 2015. The resource is continuously updating, and data extracted for this project were current to January 13, 2017. The Vanderbilt University Institutional Review Board approved this project (IRB #151116).

### Gold Standard PCS Definition

PCS was defined as one or more post-concussion symptoms (Supplementary Table 1) that persisted beyond 14 days of an mTBI. The TBI was considered moderate/severe, and subsequently excluded, if it was: (i) penetrating; (ii) accompanied by a neurosurgical procedure including hemorrhage evacuation, decompression, or invasive intracranial pressure monitoring within 7 days; (iii) associated with a GCS score <13; or (iv) accompanied by a hospitalization >5 days. A strict concussion definition was also considered, in which the TBI was ineligible in the presence of positive brain imaging.^5^ Additional exclusion criteria were <5 years of age, or any confounding neurological disease diagnosed before the mTBI (brain tumor, stroke, seizures, meningitis, intracranial abscess, multiple sclerosis, Alzheimer disease, Parkinson disease, or other cerebral degeneration).

### Sample Selection

#### (i) Natural Language Processing (NLP) Algorithm

We developed a multi-step NLP algorithm that leveraged contextual information in clinical documents to diagnose PCS cases and controls. The algorithm identified keywords for “post-concussion syndrome” and PCS symptoms (Supplementary Table 1) from the narrative text of clinical notes and problem lists using regular expression logic built into Netezza Structured Query Language. Keyword misspellings were defined a-priori (e.g., “postconcussion syndrome”), and mentions were excluded if a negation phrase occurred within 15 characters before a keyword (e.g., “no evidence of post-concussion syndrome”). Negation phrases were not expanded to characters after the keyword, as this was more likely to capture a negation phrase not associated with the original keyword. Text was not vectorized, and there were no filters on problem lists, since at VUMC, these are completed anew by providers at each clinical encounter.

The NLP algorithm was augmented by filters on coded data, and proceeded in five steps (Figure 1). In step 1, patients with mTBI were identified using a list of ICD-9 and ICD-10 mTBI codes (Supplementary Table 2). In step 2, we excluded patients with evidence of severe TBI, defined as *any* TBI code (Supplementary Table 3) followed by a neurosurgical procedure code (ICD-9 01* and 02*; ICD-10 00*, 0N*, R40.242, R40.243; Current Procedural Terminology 61000-62258) within 7 days. These neurosurgical procedure codes were previously found to be 97% sensitive and 94% specific for severe TBI.^28^ In step 3, we identified the most recent eligible mTBI using discharge windows as a proxy for TBI severity, and excluded mTBI codes that fell within a discharge window of >5 days. If a patient had multiple mTBI codes and the most recent code fell within a discharge window >5 days, the algorithm iterated over the second most recent mTBI code in the chart, checked whether it fell within a discharge window >5 days, and so on until an eligible mTBI code was found, or no previous mTBI codes remained. If a patient had no eligible mTBI codes, they were excluded. In step 4, patients meeting the following criteria were excluded: <5 years of age at the mTBI; a moderate or severe TBI (defined as any TBI code occurring in a discharge window >5 days) within 365 days of the mTBI; or a history of confounding neurological disease diagnosed before the mTBI (Supplementary Table 4).

**Figure 1.**
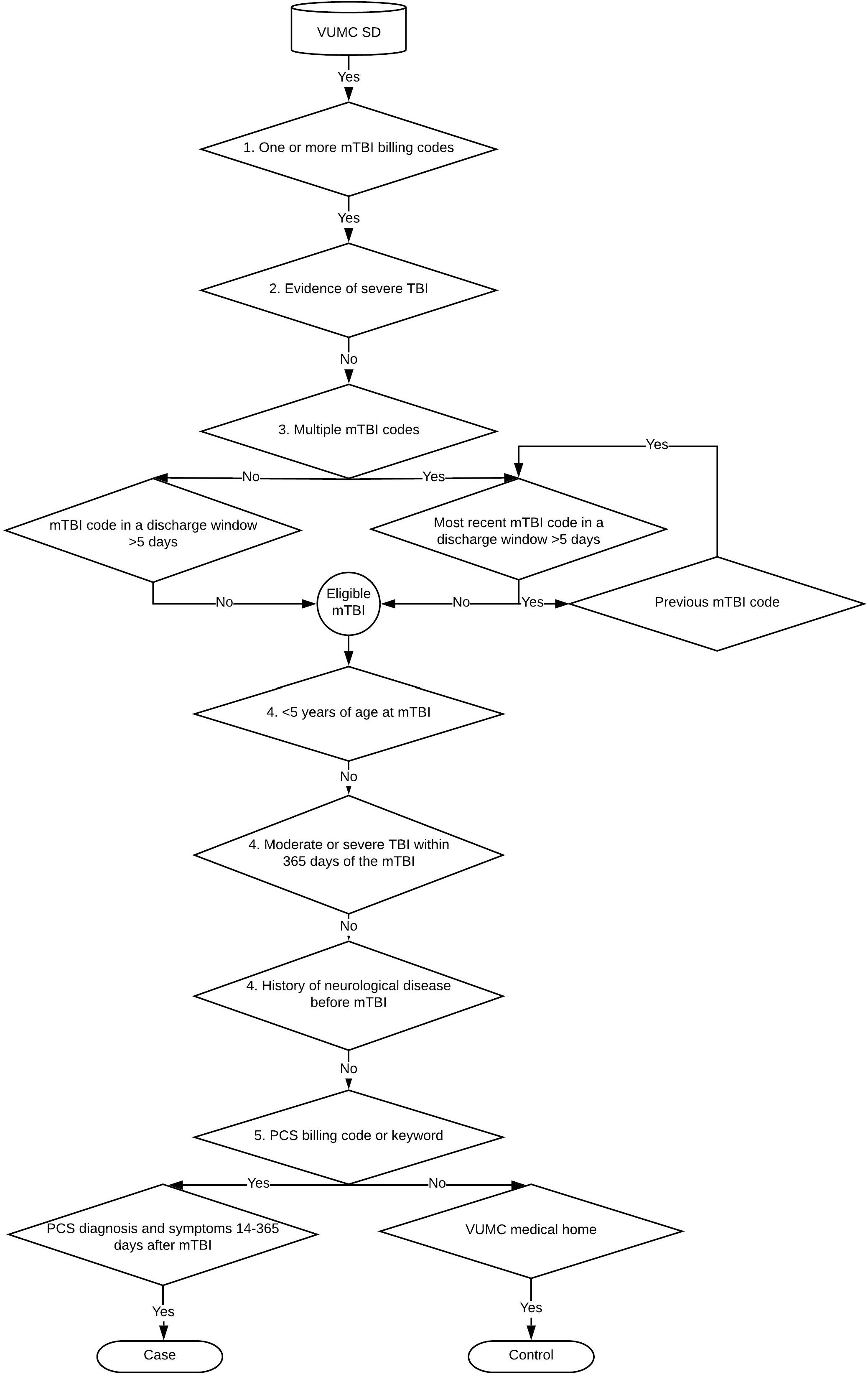
Inclusion and exclusion criteria used by the natural language processing algorithm to select mild traumatic brain injury (mTBI) patients from the Vanderbilt University Medical Center Synthetic Derivative (VUMC SD), with and without subsequent post-concussion syndrome (PCS).

In step 5, patients were classified as cases and controls based on the occurrence of PCS billing codes (ICD-9 310.2, ICD-10 F07.81) or keywords. Controls had no evidence of PCS anywhere in their record, and additionally had to have visited VUMC as an outpatient at least once in the 365 days before and once in the 365 days after the mTBI (i.e., using VUMC as their “medical home”). Cases had to have a PCS billing code or keyword on the same day as at least one PCS symptom billing code or keyword (Supplementary Table 1), and the PCS code or keyword needed to occur at least 14 days and up to 365 days after the mTBI. Patients identified by this algorithm are hereafter called NLP cases and NLP controls.

#### (ii) Coded Algorithm

We selected a second case group entirely independent of the NLP algorithm, based on only filters applied to coded data. The algorithm filters were deliberately minimal, to evaluate the performance of a simple case ascertainment scheme that maximized the number of included patients. Cases were required to have two or more instances of the PCS billing code (ICD-9 310.2, ICD-10 F07.81) on different days, a widely adopted standard for defining cases in EHR research,^25, 29^ and we excluded patients with any neurosurgical procedure code in their record (ICD-9 01* and 02*; ICD-10 00*, 0N*, R40.242, R40.243; Current Procedural Terminology 61000-62258), who were <5 years of age at their first PCS code, and those with confounding neurological histories diagnosed before their first PCS billing code. Patients identified by this algorithm are hereafter called coded cases.

### Variable Definitions

Structured data were used to describe the injury and symptom characteristics of patients identified by the NLP and coded algorithms. Variable definitions were based only on structured data, for ease of comparability across the NLP and coded algorithm patient groups. TBI morbidity group was derived by mapping ICD codes to categories of skull fracture, contusion, hemorrhage, concussion, and other/unspecified (Supplementary Table 3). Brain imaging was captured by ICD billing and procedure codes (Supplementary Table 5). TBI cause was assigned by mapping ICD external cause codes to categories of motor vehicle collision, fall, struck by and against, and assault (Supplementary Table 6). Finally, data on PCS symptoms were extracted from ICD codes (Supplementary Table 1).

### Manual Review

The validity of both algorithms was evaluated by manually reviewing the clinical documents of 50 NLP cases, 50 NLP controls, and 50 coded cases. Records were randomly selected by JD, ensuring that the NLP and coded case groups were non-overlapping. We extracted data on TBI severity, care sought, cause of injury, and subsequent PCS symptoms. A REDCap data collection instrument was piloted on 20 records, 10 of which were reviewed in triplicate (JD, PK, and AYK or SLZ), with high inter-rater agreement (Cohen’s kappa=0.80). The remaining 140 records were reviewed by PK, with all disagreements resolved by consensus. Reviewers were not blinded to case status.

### Statistical Analyses

The NPV of the NLP algorithm was the proportion of manually-reviewed controls with no PCS symptoms persisting beyond 14 days of an eligible mTBI or concussion. The PPV of the NLP and coded algorithms were the proportion of manually-reviewed cases that met the broad mTBI and strict concussion definitions of PCS. In evaluating records, we excluded first on the presence or absence of PCS symptoms, and second on TBI severity. We chose this order because PCS is ultimately defined by the persistence of symptoms, and we wanted to count the number of potential cases who were excluded based on TBI severity.

In addition to predictive values, EHR diagnostic algorithms can be validated by replicating known epidemiological associations.^25, 26^ We therefore compared each case group to controls on age, sex, and pre-morbid diagnoses that have previously been associated with PCS risk: ADHD, anxiety, depression, learning disability, migraine, and post-traumatic stress disorder (PTSD). Diagnoses were captured by billing codes (Supplementary Table 7), and we only considered those that occurred before the eligible mTBI in NLP patients, and before the first PCS code in coded patients. An age difference was detected by a t-test while categorical variables were evaluated by chi-square tests. Statistical tests were two-sided, and all analyses were conducted in R version 3.3.^31^

## Results

### Sample Characteristics

The NLP algorithm identified 28,114 patients with one or more mTBI billing codes, 26,857 with an eligible mTBI, and 24,255 potential cases and controls after excluding patients <5 years of age at the eligible mTBI, those with evidence of a moderate/severe TBI following the mTBI, and those with neurological disease preceding the eligible mTBI. Of these, 507 cases and 10,857 controls met our inclusion criteria. Cases were on average two years younger than controls (Table 1), and were more likely to be female. More than half of mTBI codes in cases and controls mapped to the “Concussion / loss of consciousness” TBI morbidity group, while the remainder mostly fell into the “Other and unspecified head injury” category. Details of the eligible TBI were more complete among controls than in cases: brain imaging was more common in controls (52.6% vs. 31.0%), as was documentation of the cause of injury (recorded in 65.7% of controls vs. 35.7% of cases), suggesting that controls were more likely than cases to have sought care at VUMC for the eligible mTBI, and that cases were only coming to VUMC and being assigned a head injury code at follow-up. Among those with a documented cause of injury, motor vehicle collision was the most common cause in both cases and controls. Nearly one fifth of patients were identified by PCS keywords as opposed to billing codes, and only 46.4% had two or more PCS billing codes. By definition, all cases had at least one PCS symptom keyword or billing code, but we used only billing code data to compare PCS symptoms across cases and controls, so comparisons could also be made with coded cases. Memory difficulties (18.1%), dizziness (17.9%), and headache (15.2%) were the most common PCS symptoms captured by billing codes in the year following the mTBI in cases, and these were much more common in cases than in controls.

**Table 1.** Characteristics of patients identified by the NLP and coded algorithms.

|  | NLP controls<br>(N=10857) | NLP cases<br>(N=507) | Coded cases<br>(N=1142) |
| --- | --- | --- | --- |
| Female gender, N (%) | 4259 (39.2) | 257 (50.7%) | 571 (50.0) |
| Age, mean (SD)<sup>a</sup> | 30.29 (18.54) | 28.25 (17.15) | 34.09 (18.36) |
| Pediatric (<18 years of age), N (%)<sup>a</sup> | 3842 (35.4) | 222 (43.8) | 294 (25.7) |
| TBI code and morbidity group, N (%)<sup>b</sup> | 10857 (100) | 507 (100) | 706 (61.8) |
| Skull fracture | 13 (0.1) | 1 (0.2) | 94 (8.2) |
| Contusion | 0 (0) | 0 (0) | 56 (4.9) |
| Hemorrhage | 0 (0) | 1 (0.2) | 120 (10.5) |
| Concussion / loss of consciousness | 6349 (58.5) | 262 (51.7) | 444 (38.9) |
| Other and unspecified head injury | 4918 (45.3) | 253 (49.9) | 560 (49.0) |
| Brain imaging, N (%) | 5714 (52.6) | 157 (31.0) | 544 (47.6) |
| TBI cause, N (%) |  |  |  |
| Motor vehicle collision | 3877 (35.7) | 90 (17.8) | 246 (21.5) |
| Fall | 1156 (10.6) | 38 (7.5) | 92 (8.1) |
| Struck by and against | 684 (6.3) | 47 (9.3) | 63 (5.5) |
| Assault | 330 (3.0) | 6 (1.2) | 20 (1.8) |
| Unknown | 4810 (44.3) | 326 (64.3) | 721 (63.1) |
| Any PCS code | 0 | 420 (82.8) | 1142 (100) |
| Two or more PCS codes | 0 | 235 (46.4) | 1142 (100) |
| PCS symptom codes<sup>c</sup> |  |  |  |
| Any | 799 (7.4) | 208 (41.0) | 582 (51.0) |
| Anxiety | 121 (1.1) | 21 (4.1) | 61 (5.3) |
| Apathy | 0 | 0 | 0 (0) |
| Depression | 325 (3.0) | 37 (7.3) | 39 (3.4) |
| Concentration difficulties | 7 (0.1) | 20 (3.9) | 13 (1.1) |
| Dizziness | 232 (2.1) | 91 (17.9) | 148 (13.0) |
| Emotional lability | 0 | 0 | 0 (0) |
| Fatigue | 109 (1.0) | 19 (3.7) | 11 (1.0) |
| Headache | 102 (0.9) | 77 (15.2) | 364 (31.9) |
| Irritable | 1 (0) | 0 | 1 (0.1) |
| Impulsive | 0 | 0 | 0 (0) |
| Memory difficulties | 164 (1.5) | 92 (18.1) | 163 (14.3) |
| Sleep disturbances | 91 (0.8) | 16 (3.2) | 32 (2.8) |
| Other | 34 (0.3) | 7 (1.4) | 31 (2.7) |

N denotes number; NLP, natural language processing; PCS, post-concussion syndrome; SD, standard deviation; and TBI, traumatic brain injury.
<sup>a</sup>Age was determined at the time of the most recent eligible mTBI code in NLP patients and at the time of the first PCS billing code in coded patients.
<sup>b</sup>Some NLP patients received multiple TBI codes on the date of the qualifying event, and some coded cases had multiple TBI codes within the window of 365 days before and 30 days after the first PCS billing code. Each TBI code was mapped to TBI type categories, which is why the row counts exceed the number of individuals in each group.
<sup>c</sup>By definition, all NLP cases had a PCS symptom billing code or keyword. For comparison with coded cases, however, in this table we only report the distribution of symptom codes. Symptom codes had to have been recorded up to 365 days after the eligible mTBI in NLP cases and controls, while in coded cases, symptom codes had to have been recorded on the same day as any PCS code.

The coded algorithm identified 5039 patients with at least one PCS code and 1372 patients with at least two PCS codes. Of these, we excluded 30 who were less than five years of age, 36 who had a neurosurgical procedure code, and 164 patients with a history of neurological disease when the first PCS code was assigned, for a final sample of 1142 coded cases (266 patients were identified as cases by both algorithms). Exactly half of cases were female. A TBI code was found in 61.8% of cases, again suggesting that many PCS patients either did not seek care for the TBI, or sought care outside of VUMC. Most TBI codes mapped to the milder categories of “Concussion / loss of consciousness” and “Other and unspecified head injury”, although a small number mapped to the more severe categories of “Skull fracture” (8.2%), “Contusion” (4.9%), and “Hemorrhage” (10.5%). Nearly half of patients had brain imaging, a proportion that was much higher than that in NLP cases, but this likely reflected differences in how brain imaging was defined in both case groups (Supplementary Table 5). In NLP patients, brain imaging codes had to occur within seven days after the eligible mTBI; in coded cases, since the date of the eligible mTBI was unknown, brain imaging codes had to occur within 365 days before and 30 days after the first PCS code. The cause of injury was documented in 36.9% of coded cases, and motor vehicle collision was again the most common cause of injury. Coded cases had a median of 3.0 PCS billing codes (range 2 to 125), with a median maximum of 65.5 days between first and last code (range 1 to 6102). Thirty-two coded cases had first and last PCS codes separated by a more than 365 days, and 334 had first and last PCS codes separated by <14 days. The top three PCS symptoms were headache (31.9%), memory difficulties (14.3%), and dizziness (13.0%), and while these were much more common in coded cases than in controls, any code was still only found in 51.0% of cases.

### Algorithm Predictive Values

The NPV of the NLP algorithm was 80% (Table 2). Two controls were excluded because they had PCS symptoms during the eligible time window and thus were true cases, but they were misclassified by the algorithm because providers had not labeled the symptoms as PCS. The first patient was <18 years of age with symptoms that resolved within 4 weeks, and recovery is known to take longer in youth.^17^ The second patient was an assault victim whose physician attributed the PCS symptoms to “tension”. Four controls were excluded because the TBI was of unclear severity, despite our efforts to ensure that controls had sufficient data in their EHRs. An additional four controls were excluded because the TBI was moderate/severe, mostly by virtue of a hospitalization >5 days. Although the algorithm had excluded patients with an eligible TBI code occurring during a hospitalization >5 days, upon manual review, the algorithm-identified eligible TBI did not always correspond to the true TBI, which in these patients was associated with a lengthy hospitalization. Of the 40 remaining controls, one had positive brain imaging, corresponding to a NPV of 78% for the strict concussion definition.

**Table 2.** Negative and positive predictive values of the algorithms determined in randomly selected sample of 150 patients.

|  | NLP controls<br>(N=50) | NLP cases<br>(N=50) | Coded cases<br>(N=50) |
| --- | --- | --- | --- |
| Symptom-based exclusion criteria applied to controls |  |  |  |
| PCS symptoms persist beyond 14 days of the index TBI<br>and are first reported <365 days after the index TBI | 2 | NA | NA |
| Symptom-based exclusion criteria applied to cases |  |  |  |
| No PCS symptoms after the index TBI | NA | 2 | 4 |
| PCS symptoms resolve within 14 days of the index TBI | NA | 4 | 0 |
| PCS symptoms are first reported >365 days after the<br>index TBI | NA | 0 | 3 |
| TBI-based exclusion criteria applied to controls and<br>cases |  |  |  |
| Index TBI was of unclear severity | 4 | 0 | 1 |
| Index TBI was moderate/severe (categories are not<br>mutually exclusive) | 4 | 2 | 4 |
| Penetrating head injury | 0 | 0 | 0 |
| Brain surgery within 7 days of index TBI | 0 | 0 | 0 |
| Glasgow coma scale score $\leq 13$ | 1 | 0 | 3 |
| Hospitalization >5 days | 3 | 2 | 2 |
| Other reasons for exclusion | 0 | 1 | 0 |
| Patients remaining | 40 | 41 | 38 |
| **Predictive value** | **NPV=80%** | **PPV=82%** | **PPV=76%** |
| Positive imaging | 1 | 3 | 5 |
| Patients remaining | 39 | 38 | 33 |
| **Predictive value (strict concussion definition)** | **NPV=78%** | **PPV=76%** | **PPV=66%** |

N denotes number; NPV, negative predictive value; PCS, post-concussion syndrome; PPV, positive predictive value; and TBI, traumatic brain injury.

The PPV of the NLP algorithm was 82%, and most cases were excluded because there was no evidence of PCS symptoms after the eligible TBI (N=2) or because the symptoms resolved within 14 days of the TBI (N=4). Three of these six patients had no PCS billing codes, only keywords. One keyword was based on a self-reported diagnosis by the patient, one was included in a negation phrase that was not captured by the algorithm (“postconcussive symptoms have resolved”) and the third was an unsupported provider diagnosis. Two of the six patients had a PCS billing code, but no symptoms consistent with PCS, and the last patient was assigned a PCS billing code at a follow-up visit during which symptoms were deemed to have resolved. After removing these patients, two additional NLP cases were excluded because the TBI was moderate/severe, again identified by a hospitalization >5 days, and a single case was excluded for other reasons (a prior pituitary tumor). Restricting to cases with negative imaging reduced the PPV to 76%.

The PPV of the coded algorithm was 76% (Table 2). Reasons for exclusion were an absence of symptoms (N=4), symptoms first reported >365 days after index TBI (N=3), a preceding TBI of unclear severity (N=1), and evidence of moderate/severe TBI (N=4). Applying the strict concussion definition reduced the PPV to 66%. The coded algorithm was simple, based mainly on PCS billing codes, and the loss of 12 patients for the above reasons shows the uneven quality of the PCS billing code. Yet, the coded algorithm has potential for improvement, and is appealing for its ease of implementation and maximal subject retention. Therefore, in an exploratory analysis, we compared the characteristics of true and false positives to identify additional features that could improve case ascertainment (Table 3). Although numbers were small, true cases were less likely to have sought care at VUMC and to have received a TBI billing code, and were more likely to have at least 14 days between their first and last PCS billing codes, and to have a PCS symptom billing code. The specialization of the provider assigning the PCS code did not differ markedly between true and false positives.

**Table 3.** Characteristics of true and false positives identified by manual review of 50 coded cases.

|  | True positives<br>(N=38) | False positives<br>(N=12) |
| --- | --- | --- |
| Care sought for index TBI, N (%) |  |  |
| Yes - VUMC | 17 (45) | 7 (58) |
| Yes - Not VUMC | 9 (24) | 0 (0) |
| No | 2 (5) | 1 (8) |
| Unknown | 10 (26) | 4 (33) |
| TBI code and morbidity group, N (%)<sup>b</sup> | 23 (61) | 9 (75) |
| Skull fracture | 3 (8) | 1 (8) |
| Contusion | 1 (3) | 1 (8) |
| Hemorrhage | 4 (11) | 2 (17) |
| Concussion / loss of consciousness | 14 (37) | 6 (50) |
| Other and unspecified head injury | 21 (55) | 6 (50) |
| First and last PCS codes separated by a minimum of 14 days, N (%) | 31 (82) | 7 (58) |
| First and last PCS codes separated by a maximum of 365 days, N (%) | 1 (3%) | 0 (0) |
| PCS symptom codes - any | 26 (68) | 6 (50) |
| Specialty of provider assigning the first PCS billing code, N (%) |  |  |
| Neurology | 15 (39) | 5 (42) |
| Primary Care | 2 (5) | 0 (0) |
| Emergency Medicine | 8 (21) | 4 (33) |
| Psychiatry | 0 (0) | 0 (0) |
| Sports Medicine | 2 (5) | 0 (0) |
| Other | 6 (16) | 0 (0) |
| Unknown | 5 (13) | 3 (25) |

### Epidemiology of PCS in EHR

To complement the manual review, we next evaluated the algorithm by comparing the distribution of known PCS risk factors across algorithm-identified cases and controls. Cases identified by both approaches were more likely than controls to be female (Table 1; p=3.25x10^-7^ for NLP cases and p=2.25x10^-7^ for coded cases). The association with age, however, was discrepant across case groups; compared to controls, NLP cases were two years younger on average (p=1.58x10^-3^) while coded cases were nearly four years older (p=4.15x10^-11^). Pre-morbid anxiety, migraine, and PTSD were more frequent in both case groups versus controls (Table 4). A higher prevalence of depression in cases was also found, but the increase was only statistically significant in the larger sample of coded cases. In contrast, diagnoses with uncertain associations with PCS, ADHD and learning disabilities, were equally common in our cases and controls, suggesting that the enrichment of pre-morbid anxiety, migraine, and PTSD in our cases captured true associations as opposed to a systematic under-reporting of diagnoses in controls.

**Table 4.** Pre-injury diagnoses in controls and cases. Diagnoses (Supplementary Table 7) had to occur before the eligible mTBI in NLP patients, and before the first PCS billing code in coded patients.

| Diagnosis | NLP controls<br>(N=10857) | NLP cases<br>(N=507) | p-value vs.<br>controls | Coded cases<br>(N=1142) | p-value vs.<br>controls |
| --- | --- | --- | --- | --- | --- |
| ADHD, N (%) | 227 (2.1) | 14 (2.8) | 0.39 | 24 (2.1) | 1 |
| Anxiety, N (%) | 560 (5.2) | 39 (7.7) | 0.02 | 112 (9.8) | 1.26E-10 |
| Depression, N (%) | 581 (5.4) | 36 (7.1) | 0.11 | 101 (8.8) | 1.74E-06 |
| Learning disability, N (%) | 129 (1.2) | 10 (2.0) | 0.17 | 19 (1.7) | 0.21 |
| Migraine, N (%) | 246 (2.3) | 35 (6.9) | 1.31E-10 | 117 (10.2) | 4.16E-50 |
| PTSD, N (%) | 100 (0.9) | 14 (2.8) | 1.25E-04 | 27 (2.4) | 1.18E-05 |

ADHD denotes attention deficit hyperactive disorder; N, number; NLP, natural language processing; and PTSD, post-traumatic stress disorder.

The above exploratory analysis of the 50 manually reviewed coded cases suggested that the coded algorithm could be improved by filtering on TBI codes and PCS code density. To further investigate the effect of these filters, we additionally report the characteristics and PCS risk factors in coded cases (1) with and without a TBI billing code; (2) with and without a moderate/severe TBI billing code; and (3) with at least 14 days between first and last PCS billing codes. Coded cases with an TBI billing code were younger and more likely to be male than coded cases without an TBI billing code (mean age 31.4 vs. 38.4 years; 53.1% vs. 45.0% male; Supplementary Table 8). Among patients with TBI billing codes, those with moderate/severe as opposed to mild TBI codes were even more likely to be male (65.3% vs. 49.3%). Conversely, imposing a minimum time between first and last PCS codes reduced the proportion of males in the sample (46.2% vs. 50.0% of all coded cases) and of patients with an TBI billing code (55.6% vs. 61.8% of all coded cases). The cardinal pre-injury PCS risk factors of anxiety, migraine, and PTSD were marginally more prevalent in coded cases without an TBI billing code vs. with an TBI billing code, and in coded cases with at least 14 days between first and last PCS codes vs. all coded cases (Supplementary Table 9). The known risk factors, however, were markedly absent in the sample of coded patients with a moderate/severe TBI billing code, suggesting that these moderate/severe TBI billing codes could be useful exclusion criteria in future algorithms.

## Discussion

TBIs are sustained by more than 1 in 120 Americans annually,^1^ and adverse outcomes affect many. In an initial attempt to capture patients with mTBI and PCS from an EHR, we report moderate predictive values for both an NLP-based algorithm, and a simpler coded algorithm that used only structured data. We also show that including all algorithm-identified patients in case-control comparisons recovered the known epidemiology of PCS, suggesting that when the sample size is large, true associations can be detected despite imperfect patient classification. Female gender, prior mood disorders, and headache history are some of the strongest predictors of poor recovery after an mTBI.^11-16^ We used only EHR data to capture pre-injury medical histories and found very statistically significant associations with these variables. Our results indicate that EHR-based subject ascertainment can substantially increase sample sizes, paving the way for discoveries of novel biological determinants of recovery which could ultimately lead to early identification and intervention for patients at risk.

The NLP algorithm had the best PPV (82%), in line with that of EHR algorithms for other neuropsychiatric disorders.^24, 32, 33^ In addition, the algorithm detected patients by PCS keywords that were not captured by the coded algorithm. These keywords, however, were of questionable validity. Moreover, the number of cases captured by the NLP algorithm was less than one tenth of the patients in the SD with a single PCS billing code, and requiring coded documentation of the TBI prior to the PCS diagnosis excluded many potential cases. The algorithm was also time-consuming to develop and test, and complicated to implement. These strengths and limitations of NLP algorithms are well recognized.^24^ EHR algorithms for autism,^32^ bipolar disorder,^34^ and major depression^24^ had the best PPV (~85%) when narrative text was included, and a systematic review of 19 algorithms across a range of chronic and infectious conditions found that incorporating narrative EHR text improved case detection from 61.7% to 78.1%.^35^ Nonetheless, fewer patients have sufficient EHR data to be classified by NLP algorithms, leading to considerable reductions in sample size.^24, 36^

The coded algorithm had a lower PPV (76%), but captured twice as many cases. Diagnostic and procedure codes can be “noisy”, yet they have been used repeatedly in EHR research to validate known environmental and genetic associations, and for novel discovery.^25, 26, 36-39^ In the largest proof-of-concept experiment, 77 robust disease-gene associations were tested for replication using disease diagnoses derived from ICD-9 billing codes in 13,835 patients.^40^ Sixty-six percent of the associations replicated, again showing that large datasets can overcome imperfect case-control assignments. The coded algorithm is therefore appealing for further development since it includes many patients and recovers known PCS risk factors, even in the presence of patient misclassification.

Another strength of billing codes is that they reflect real clinical practice, and discoveries made with these data may better translate into clinical advances than discoveries made in highly ascertained samples. Real-life data are messy: physicians assigned PCS billing codes even when patients had moderate/severe TBI. We excluded these patients, but arguably they should be included in future studies of PCS since they represent the true clinical spectrum of disease. Moreover, the biology that underlies TBI recovery may be insensitive to TBI severity. Large prospective registry-based studies have found an increased incidence of adverse psychological, cognitive, emotional, and social outcomes across the TBI severity spectrum, just to a lesser extent in mTBI patients.^41-44^ Future EHR-based studies of PCS could therefore test all PCS patients, regardless of TBI severity, in sensitivity analyses.

We compared our algorithms’ performance to a gold standard PCS definition, but diagnostic criteria for PCS are contentious.^45^ The term PCS was first used in the 1930s. In the ICD-9 manual, PCS was described as a constellation of symptoms, and in the DSM-IV and ICD-10 manuals, explicit diagnostic criteria were provided.^46^ The agreement between DSM-IV and ICD-10 criteria, however, is poor,^46^ the DSM-5 has removed the PCS diagnosis entirely in favor of defining mild or major neurocognitive disorder following TBI,^30^ and few clinicians even adhere to DSM or ICD guidelines in practice.^47^ Large-scale EHR data offer new opportunities to resolve these diagnostic discrepancies by revealing how TBI patients actually cluster and recover clinically. Data-driven approaches have already uncovered novel autism spectrum disorder subgroups using the EHR,^48^ and machine learning algorithms applied to EHR data have accurately predicted suicide attempts up to two years in advance.^49^ Our study is an important first step towards applying similar tools to the EHR of TBI patients.

### Limitations and Strengths

Algorithms were developed and tested at only a single institution. Also, the PPVs were modest, but in line with the PPVs of EHR algorithms published for other psychiatric diseases.^24, 32, 34^ The NPV of the NLP algorithm, on the other hand, was lower than expected. Insufficient data to classify TBI severity was a leading reason for excluding controls, emphasizing the challenges of using EHR data for research when care is fractured across providers. In addition, our study was only designed to determine the algorithms’ predictive values, not their sensitivity or specificity; calculation of these parameters would require a gold standard case-control set. Nonetheless, this study demonstrates the feasibility of ascertaining large samples of PCS cases and controls from the EHR, and is a valuable starting place for future algorithm development efforts.

### Recommendations for Future EHR-Based Algorithms of PCS

Our analyses revealed the advantages and disadvantages of different approaches for identifying PCS cases and controls from EHRs. Recommendations for future algorithms are: (1) Select cases by PCS billing codes that meet a pre-specified code density threshold. This criterion will help remove cases who were assigned PCS billing codes at the time of injury, without subsequent evidence of symptom persistence. Do not require a TBI billing code, as this enriches the sample for patients who seek care only for the index TBI and not for the persistence of symptoms, and filter judiciously on PCS symptom codes to avoid removing too many potential cases. (2) Exclude cases with moderate/severe TBI billing codes. These codes mostly captured cases who sought care for the index TBI, and not for ongoing PCS symptoms. (3) Select controls with mTBI billing codes, and filter those with neurosurgical procedure codes, moderate/severe TBI billing codes, lengthy hospitalizations, and insufficient pre- and post-injury data. Although these recommendations violate epidemiological first principles (a case is an eligible control, were it not for disease diagnosis), requiring cases to have EHR documentation of the TBI (as in our NLP algorithm) was too restrictive. Instead, we suggest imposing strict TBI severity filters on controls to minimize potential bias caused by controls having more severe TBI or greater health-seeking behavior relative to cases.

## Conclusions

EHR hold great promise for understanding TBI sequelae. We show that EHR are valid for TBI research, and set the stage for more extensive data linkages, including to biorepositories with genetic data. Two strategies were employed in the current study to ascertain patients with mTBI and PCS from the EHR, and each strategy had tradeoffs. On balance, however, the coded algorithm had a reasonable PPV, recovered known PCS associations and maximized the number of included patients. No single approach will be perfect, and future studies should continue to compare different diagnostic algorithms. EHRs are a vast, growing resource available to many independent research groups, and the use of EHR-linked biobanks in TBI research will have far-reaching public health benefits.

## Acknowledgements

The authors thank Lea K. Davis and Rebecca T. Levinson for comments on earlier iterations of the manuscript, and Doug Conway for help implementing the algorithms and navigating access to the EHR data.

Jessica Dennis is supported by the Canadian Institutes of Health Research (award MFE-142936). Support for Nancy J. Cox was provided by R01 MH113362, U54MD010722, and U01HG009086. This project was supported by a Vanderbilt Institute for Clinical and Translational Research micro-grant (VR20299) and was conducted in part using the resources of the Advanced Computing Center for Research and Education at Vanderbilt University, Nashville, TN. The datasets used for this project were obtained from Vanderbilt University Medical Center’s Synthetic Derivative, which is supported by numerous sources: institutional funding, private agencies, and federal grants. These include the NIH funded Shared Instrumentation Grant S10RR025141; and CTSA grants UL1TR002243, UL1TR000445, and UL1RR024975. The REDCap tool used in this study was supported by grants UL1 TR000445 from NCATS/NIH.

## Author Disclosure Statement

Gary Solomon is a consultant for the Nashville Predators, Tennessee Titans, and the athletic departments of Tennessee Tech University and the University of Tennessee, fees paid to institution. He is also a consultant to the National Football League Department of Health and Safety.

The other authors have no competing financial interests.

